# Adaptive admixture at *ACKR1* (the Duffy locus) may have shaped *Plasmodium vivax* prevalence in Oman

**DOI:** 10.1101/2024.03.06.583766

**Authors:** Paige E. Haffener, Arwa Z. Al-Riyami, Shoaib Al-Zadjali, George B. J. Busby, Sulaiman Al Mahdhuri, Mohammed Al-Rawahi, Saif Al Hosni, Ali Al Marhoobi, Ammar Al Sheriyani, Ellen M. Leffler

## Abstract

Malaria in humans is largely caused by two divergent species of *Plasmodium* parasites, *P. vivax* and *P. falciparum*, both of which have driven the spread of protective alleles in human populations. Notably, an erythrocyte-specific Duffy null allele (Fy^ES^) confers resistance to *P. vivax* malaria and has been identified as a target of strong, recent positive selection in multiple African admixed populations. Here, we evaluate evidence for selection via adaptive admixture in Oman, where compared to neighboring countries, *P. vivax* has recently been less common. Genetic ancestry inference using whole genome sequence data from 100 Omanis suggests 9.8% (95% CI: 7.3-12.2%) of their genetic ancestry is shared with east Africa. At the Duffy locus, we find a high frequency of Fy^ES^ and an increase to 76% African ancestry. Comparing with blood group serology for the same individuals, we identify an additional Duffy-null allele that is rare but present across multiple Arabian Peninsula (AP) populations. Finally, we estimate the selection coefficient at Fy^ES^ as 0.031 (95% CI: 0.029-0.034) with likely introduction at least 59 generations ago, older than estimates in other African admixed populations. Although we also observe higher frequency of some *P. falciparum*-protective alleles in Oman than in other AP populations, African ancestry is not enriched indicating a lack of evidence for adaptive admixture driven by *P. falciparum* selective pressure. Together, our analyses suggest that Oman’s long history with east African populations resulted in early introduction and selection for Duffy null alleles and may have influenced the prevalence of *P. vivax* in the region.

## Introduction

Infectious pathogens are a major selective pressure across the tree of life, driving adaptations in host species that in turn affect pathogen distribution and evolution. A classic case of host-pathogen arms race exists between humans and malaria parasites and is exemplified between *Plasmodium vivax* and the Duffy antigen receptor for chemokines (DARC), which is a critical receptor for the invasion of red blood cells by *P. vivax*^1^. One of the strongest selection signatures in the human genome occurs at the gene encoding DARC (*ACKR1,* atypical chemokine receptor 1), which also determines the human Duffy blood group. The erythrocyte null, or silent (ES), allele Fy^ES^ (−67 T>C rs2814778) prevents DARC from being expressed on red blood cells^2^ and is associated with protection against invasion by *P. vivax*^1^. It is the most highly differentiated single nucleotide polymorphism (SNP) between European and African populations in the 1000 Genomes Phase 3 dataset^3^. This extreme differentiation is inferred to be the result of a selective sweep 42,000 years ago (post-dating the out of Africa expansion) leading to near fixation of Fy^ES^ and the near absence of *P. vivax* from sub-Saharan Africa^3^, a hypothesis established roughly 50 years ago^1,4–6^.

The Fy^ES^ allele has also been reported in multiple populations with African admixture^7–11^ at higher frequency than expected under models of genetic drift, suggesting repeated convergent positive selection, i.e. adaptive admixture. In the African-admixed populations of Madagascar and Cabo Verde, the strength of selection at the Fy^ES^ allele has been estimated between 0.07 and 0.2^7–9^, comparable to estimates of selection coefficients at malaria-associated loci such as the HbS, G6PD A-, and G6PD Med alleles^12,13^. Although selection coefficients were not estimated, positive selection at the *ACKR1* gene has similarly been inferred through the observation of significant excesses of African ancestry in Pakistani, Iranian, Saudi Arabian, Yemeni, Emirati, and Omani populations^10,11^. Selection for the Fy^ES^ allele in each of these populations is thought to be quite recent, occurring within the last 500 to 1,000 years with introduction of the allele attributed to large movements of peoples through the Indian Ocean slave trade and colonization^9–11,14^.

The Arabian Peninsula (AP) has served as a bridge connecting African and Eurasian populations for thousands of years. While the earliest evidence places some level of contact between AP and African and Eurasian populations as early as 10,000 years ago^15^, most African admixture into AP populations is thought to have occurred in the last 1,000 years, coinciding with the southern AP population maritime commerce and Indian Ocean slave trade^11,15,16^. In connection, southern AP populations have been shown to have more sub-Saharan African admixture than AP populations to the north (Supplementary Figure 1). For instance, in Yemen and Oman, countries on the southern border of the AP, estimates of sub-Saharan admixture are approximately 10% whereas in the United Arab Emirates (UAE) estimates range from 3 - 10% and from 3 - 7% in Saudi Arabia^11,16^. In northern populations from Iraq, Jordan, and Syria, estimates are much lower, ranging from 0 - 2%^16^. Oman in particular has sustained recurrent historical connections to East Africa including migrations in the 7^th^ and 13^th^ centuries and establishment of the Omani Empire along the Eastern African coast dating to the 17^th^ century (Supplementary Table 1).

Putative positive selection for the Fy^ES^ allele following African admixture in the AP is also supported by a relatively high frequency of the Duffy negative phenotype, defined as the serological absence of Duffy antigens, across the AP^17–21^, most notably in Oman^22^. While the Duffy negative phenotype has been linked to *P. vivax* resistance, little documentation exists regarding the historical prevalence of malaria in Oman or other AP countries. Records indicate that malaria has persisted in Oman since the earliest documentation in the 1900s to 2013 when the World Health Organization (WHO) determined Oman to be malaria free due to successful control efforts^23^. Although data is sparse, prior to elimination efforts less than 5% of cases in Oman were caused by *P. vivax*^23–25^ in contrast to approximately 50% in Saudi Arabia and 20-30% in the UAE^25^. Recent data from the early stages of eradication efforts, as indicated in the WHO’s annual World Malaria Reports, also support this pattern. For example, in 2003, *P. vivax* accounted for 80% and 100% of malaria cases in Iran and Iraq, respectively^26^. In contrast, Saudi Arabia had 27% of cases attributed to *P. vivax,* while Yemen had only 2% with the remainder due to *P. falciparum*.

We hypothesize that historical African admixture and selection for Duffy null alleles may have led to the high prevalence of the Duffy negative phenotype and exclusion of *P. vivax* from Oman, akin to the hypothesis explaining the limited presence of *P. vivax* in sub-Saharan Africa. Here, we analyze Duffy serology and whole genome sequences from 100 Omanis, increasing from only three genomes previously sequenced^16^, to investigate the presence of null alleles in *ACKR1* that may have been the target of selection on the locus. We further use genetic ancestry inference to confirm the origin of the Fy^ES^ allele in Oman as well as to assess the strength of selection and timing of introduction.

## Methods

### Sample Collection

Donors were recruited at the Sultan Qaboos University Hospital (SQUH) blood bank between January and June 2020. Randomly-selected, healthy male and female blood donors ranging in age from 18 – 60 were interviewed by a research member and were consented for enrollment. Any non-Omani donors or any donor with history of recent transfusion within 6 months’ period of the enrollment were excluded from the study. The SQUH blood bank in Muscat draws individuals from all major regions of Oman and represents a cross-section of the Omani population. Two EDTA samples (Ethylenediaminetetraacetic acid, an anticoagulant) were collected from each donor upon consenting for enrollment. The study was approved by the Medical Research Ethics committee of Sultan Qaboos University.

### Blood Bank Methods

Red blood cell phenotyping was performed at SQUH Blood bank on freshly drawn samples within 24 hours of collection. Phenotyping was performed as per the manufacture instructions (BioRad©, Cressier Switzerland) and as previously published^22^. In brief, reactions were graded from 0 to 4 based on the strength of agglutination. A grade 0 or 1 reaction was regarded as negative. Grades 2,3 and 4 reactions were regarded as positive. Known positive and negative samples were included as internal controls. For this study, we report results for the Duffy blood group system (Fy^a^,Fy^b^ antigens in Antigen Profile III).

DNA extraction was performed at the Department of Hematology, SQUH. Whole blood was collected with EDTA as an anticoagulant, and genomic DNA was extracted using the QIAamp DNA Blood mini kit (Qiagen, Inc., Valencia, CA) as per the manufacturer’s instructions. After extraction, samples were saved at −20 degrees until shipped at room temperature using a certified courier to the University of Utah.

### Whole Genome Sequencing

Short read (150bp) whole genome sequencing was performed at the Huntsman Cancer Institute High-Throughput Genomics Shared Resource at the University of Utah on the Illumina Novaseq 6000. Samples were multiplexed and sequenced to an average coverage of 16X.

### Read mapping and variant calling

Raw sequencing reads were aligned to GRCh38 (ftp://ftp.ncbi.nlm.nih.gov/genomes/all/GCA/000/001/405/GCA_000001405.15_GRCh38/seqs_for_alignment_pipelines.ucsc_ids/GCA_000001405.15_GRCh38_no_alt_plus_hs38d1_analysis_set.fna.gz; downloaded Dec. 07, 2020) using BWA-MEM^27^. PCR duplicates were removed with samtools^28^. Unmapped reads, duplicates, non-primary alignments, and reads with a MAPQ < 20 were filtered out for coverage calculations using ‘bedtools genomecov’^29^. Sample sex was confirmed by comparing the coverage of chromosome 7 to the coverage of chromosome X. Lane-level read groups were added and BAM files were merged with samtools to generate a sample-level BAM file for each individual. Variants were called using Sentieon^30^ to implement the GATK best practices protocol^31^. We checked for close relatives using king v1.4^32^, but found no first or second degree relatives and kept all 100 samples for analysis.

### Global Ancestry Inference and Principal Component Analysis (PCA)

The Omani variants (in variant call format, VCF) were merged with the 1000 Genomes Phase 3 VCF files (http://ftp.1000genomes.ebi.ac.uk/vol1/ftp/data_collections/1000_genomes_project/release/20190312_biallelic_SNV_and_INDEL/) matching by position and allele using bcftools^33^ and plink2^34^. Preparation for PCA and admixture analyses included LD pruning according to the ADMIXTURE^35^ manual, filtering out variants with MAF < 0.05, and excluding AMR samples using plink2. PCA analysis was done using Eigensoft smartpca version 13050^36,37^. ADMIXTURE v 1.3.0 was run using K = 11 source populations, as determined by cross-validation, and results were plotted with AncestryPainter v5.

### Local Ancestry Inference with RFMix v2

The Omani genotypes were phased using Eagle v 2.4.1^38^ and the genetic map provided for GRCh38. We used the high coverage 1000 Genomes GRCh38 for the reference genotypes. RFMix v2^39^ was run using 100 EM iterations and specifying 10 generations from time of admixture. To calculate excess ancestry, we summed the number of segments in the genome classified as each ancestry and divided by the total number of chromosomes and plotted the sums against the chromosomal positions. For each chromosome, the genome-wide mean and variance was determined for each ancestry excluding the chromosome under consideration.

### Admixture Dating

To estimate the date of admixture using MALDER, we generated five random samples of 750,000 biallelic SNVs from the merged VCF by randomly sorting all VCF positions and creating a subset VCF with the first 750k variants from each random sorting event. The randomly sampled VCFs were converted to EIGENSTRAT format using a pipeline of plink and Eigensoft v 6.0.1. These files were used as input for MALDER^40^ to infer admixture dating events via linkage disequilibrium decay using 1000 Genomes populations Esan in Nigeria (ESN), British in England and Scotland (GBR), and Sri Lankan Tamil in the UK (STU) as reference populations. To infer the date of admixture with Multiwaver2.0^41^, we obtained ancestry tract lengths for reference populations – Luhya in Webuye, Kenya (LWK) and Other (Toscani in Italia (TSI) + STU) – from the RFMix v2 local ancestry output. Multiwaver2.0 was run using 100 bootstrap replicates and the ‘MixMode’ argument to infer the population model best fit for the data.

### Inferring the selection coefficient and initial African ancestry

We simulated local ancestry and positive selection following the introduction of the Duffy negative allele using SLiMv4^42–45^, running 1000 simulations for each demographic model, and basing our models off the continuous and single pulse simulation models from Hamid et al.^9^. We set the number of generations to 10 to simulate an early pulse similar to the admixture date inferred by MALDER, 59 to match the admixture date inferred by MultiWaver, and 75 to test an older introduction. The effective population size (*Ne*) was set to 15,000^16^, and an exponential growth rate of 5%. We set the migration rate, or starting African (*p1*) ancestry proportion (*mig*), to be sampled from a uniform distribution of 0.10 – 0.90. We set the selection coefficient (*s*) to also be sampled from a uniform distribution of 0.0 – 0.20. Using the provided python scripts from SLiM recipe 17.5, we then calculated five summary statistics for each simulation from the local ancestry tree output files. These were: African ancestry proportion at Duffy, variance in global African ancestry, median tract length at Duffy, mean tract length at Duffy, and variance in tract length at Duffy.

The same five summary statistics were calculated for the Omanis using output from RFMix v2 and ADMIXTURE. We obtained the sampled *s* and *mig* values for each of the 1000 simulations and used them to estimate *s* and *mig* in Oman with Approximate Bayesian Computation (R package abc)^46^. The goodness-of-fit of the summary statistics was determined with ‘gfit’, and the prediction error of the s and mig parameters was calculated using ‘cv4abc’ with 100-fold cross validation and tolerance values of 0.5%, 1.0%, and 5.0% on the 1000 sets of summary statistics. Using the ‘neuralnet’ method and the tolerance with the lowest prediction error, 5.0%, we inferred parameter values for *s* and *mig* with ‘abc’.

### Copy number calling at the *GYP* gene cluster

We inferred copy number of the *GYP* gene cluster on chr4 using a HMM previously developed for this region^47^ and available on github (https://github.com/malariagen/glycophorin_cnvs/blob/master/call_cnvs/README.md). Following the previous approach, we excluded sites with low mappability (mean CRG mappability <0.9 for all 100-mers overlapping the site) and excluded windows (window size of 1600bp) with < 25% mappable sites. We also included coverage for HG02554 from 1000 Genomes, as it is known to carry the Dantu structural variant, to facilitate comparison to the Omani data.

### Sanger sequencing of the ACKR1 frameshift allele

We used the ThermoFisher primer pair Hs00587748_CE for sequencing the region of the *ACKR1* locus containing the identified 2bp frameshift deletion allele. These primers amplify a 234bp product (chr1:159,205,472-159,205,705). We followed the Phusion Flash Master Mix protocol and calculated the annealing temperatures using the calculator provided by ThermoFisher. We used 34 cycles and an annealing temperature of 64.3°C. The recommend amount of DNA to amplify for a product of 234bp is between 50ng and 75ng. We amplified 50ng of DNA for all Omani samples except for the additional samples OM11, OM21, OM29, and OM42 for which we used 75ng because DNA yield was too low for sequencing when amplifying 50ng with these samples. 30uL of PCR product was mixed with 6uL of 6X TriTrack DNA loading dye (Catalog #R1161) and 5uL of GeneRuler 50bp DNA Ladder (Catalog #SM0373) were loaded on a 1% gel with Ethidium Bromide and run at 57V for 2.5 hours. Purified DNA was recovered using the Zymoclean Gel DNA Recovery Kit (Catalog #11-300C). Samples were submitted to the University of Utah DNA Sequencing Core following the guidelines for sample submission based on PCR product size. We used 4Peaks^48^ to analyze the Sanger sequencing traces, a tool that includes visualization of the traces, outputs the sequence, and reports sequence quality.

## Results

### Genomic evidence for recent East African and South Asian admixture in Oman

We performed Illumina whole genome sequencing and variant calling with GATK on 100 healthy Omani blood donors (mean coverage=16X; 25.8 million SNPs and short indels; Supplementary Table 2). For comparison with global populations, we merged this dataset with the 1000 Genomes phase 3 variant calls^49,50^, resulting in an intersection set of approximately 18 million variants (Supplementary Table 2).

Global genetic ancestry inference with ADMIXTURE^35^ across all 2604 individuals identified a distinct ancestry component in Oman (mean proportion 79%, 95% confidence interval (CI) 75-83%). This component is absent across 1000 Genomes populations (with the exception of the Toscani in Italia (TSI) mean proportion 5.3%, 95% CI 5.1-5.5%) consistent with global ancestry analyses of other Middle Eastern populations when analyzed together with the 1000 Genomes dataset, which has no representation from the Middle East (Figure 1A)^11,51–54^. The second largest ancestry component in the Omanis (mean proportion 9%, 95% CI 7.3-12.2%) is shared specifically with the Luhya in Webuye, Kenya population (LWK), the only east African population in the dataset. The Omanis also shared ancestry components common to both east and west African populations (mean proportion 6.2%, 95% CI 4.43-7.98%) and South Asian populations (mean proportion 2.3%, 95% CI 1.2-3.3%).

**Figure 1.**
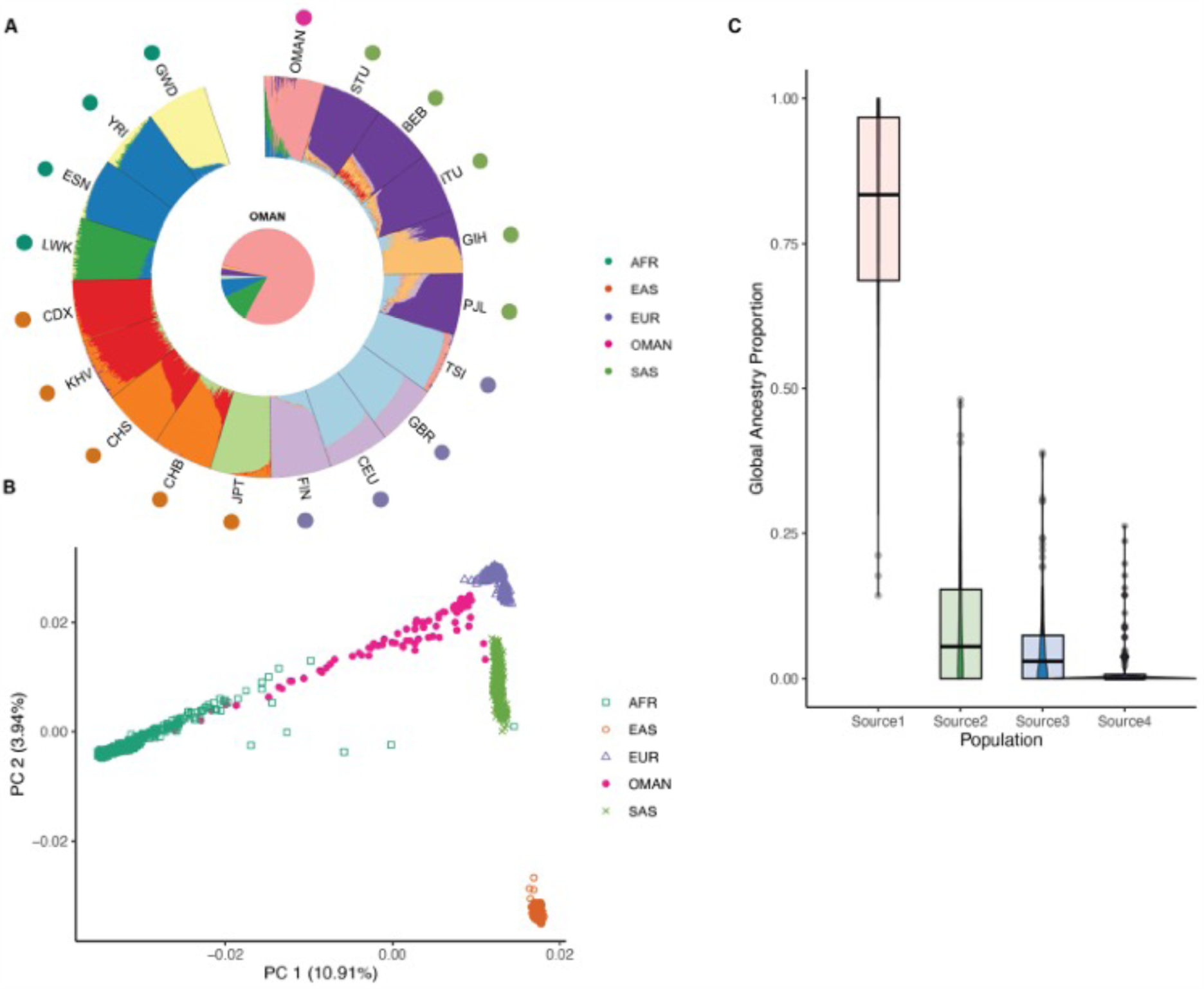
Inference of global ancestry and population structure. A) Global admixture inferred using ADMIXTURE with k = 11 source populations. Populations included are from the 1000 Genomes AFR (African), EAS (East Asian), EUR (European), and SAS (South Asian) super populations as well as the 100 Omani individuals. The pie chart in the middle shows the overall proportion of each source population across the 100 Omani individuals. B) The first two principal components from PCA of genome-wide SNP data with Omani individuals projected onto the principal components computed from the 1000 Genomes. C) Variation in ancestry from the four largest contributing source populations across Omani individuals. Colors are the same as in A, with source 1 most common in Oman, source 2 most common in east Africa, source 3 in both west and east Africa, and source 4 in south Asia. Population abbreviations are: GWD: Gambian in Western Division, Mandinka. YRI: Yoruba in Ibadan, Nigeria. ESN: Esan in Nigeria. LWK: Luhya in Webuye, Kenya. CDX: Chinese Dai in Xishuangbanna, China. KHV: Kinh in Ho Chi Minh City, Vietnam. CHS: Southern Han Chinese. CHB: Han Chinese in Beijing, China. JPT: Japanese in Tokyo, Japan. FIN: Finnish in Finland. CEU: Utah residents (CEPH) with Northern and Western European ancestry. GBR: British in England and Scotland. TSI: Toscani in Italia. PJL: Punjabi in Lahore, Pakistan. GIH: Gujarati Indians in Houston, TX, USA. ITU: Indian Telugu in the UK. BEB: Bengali in Bangladesh. STU: Sri Lankan Tamil in the UK.

Local ancestry inference using RFMixv2^39^ with representative 1000 Genomes populations from east Africa (LWK), west Africa (Esan in Nigeria, ESN), south Asia (Sri Lankan Tamil in the UK, STU) and a proxy Middle Eastern population (TSI) yielded similar African admixture proportions. RFMixv2 inferred an average of 11.09% LWK admixture (95% CI 11.05-11.13%), 5.89% ESN admixture (95% CI 5.86-5.92%), and 0.11% STU admixture (95% CI 0.107 - 0.115%) in the Omanis. In further support of both admixture inference results, PCA placed the Omanis along a cline with similarity to both African and South Asian populations (Figure 1B). We also observed wide variability in admixture across individuals (Figure 1C), particularly in global African admixture (0.001%-48.1% east African component), suggesting some recent or ongoing admixture in Oman^55^.

Genetic ancestry in Omanis shared with East African and South Asian populations is consistent with the documented strong and sustained historical trade connections across the Indian Ocean^56,57^. To estimate the timing of historical admixture with African and South Asian populations, we performed admixture dating with MALDER^58^, which can estimate up to two discrete admixture pulses based on the decay of admixture-introduced linkage disequilibrium (LD). Using representative 1000 Genomes populations from Africa (ESN), Europe (British in England and Scotland, GBR), and South Asia (STU) as source populations, MALDER inferred a single pulse of African admixture in the Omanis occurring approximately nine generations ago (Table 1).

**Table 1.**
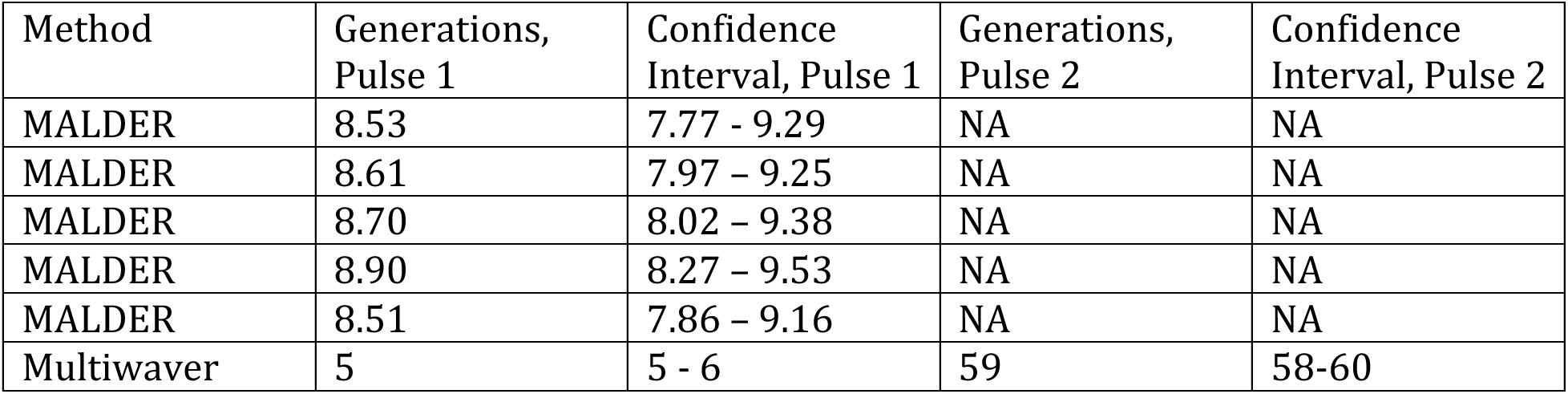
Dates of African admixture in Oman inferred by MALDER and Multiwaver2.0. Each MALDER row represents an independent random sampling of 750,000 bi-allelic SNPs.

| Method | Generations, Pulse 1 | Confidence Interval, Pulse 1 | Generations, Pulse 2 | Confidence Interval, Pulse 2 |
| --- | --- | --- | --- | --- |
| MALDER | 8.53 | 7.77 - 9.29 | NA | NA |
| MALDER | 8.61 | 7.97 - 9.25 | NA | NA |
| MALDER | 8.70 | 8.02 - 9.38 | NA | NA |
| MALDER | 8.90 | 8.27 - 9.53 | NA | NA |
| MALDER | 8.51 | 7.86 - 9.16 | NA | NA |
| Multiwaver | 5 | 5 - 6 | 59 | 58-60 |

However, one or two discrete pulse models of admixture may not be a good fit, as multiple periods of increased connectivity in the region have been described based on historical records (Supplementary Table 1) and MALDER can fail to differentiate between ancestry pulses from similar ancestry groups. To assess additional scenarios, we also inferred admixture dates using MultiWaver2^41^, which uses the distribution of tract lengths rather than LD decay and is able to detect more complex population histories, evaluating continuous, multi-pulse, and single-pulse models^59^. Using this approach, we infer a two-pulse model of African admixture occurring 5 and 59 generations ago (bootstrap support = 100%, Table 1).

### An excess of African ancestry at ACKR1 indicates strong positive selection

Admixture provides an opportunity for advantageous alleles to spread from one population to another. This process, known as adaptive admixture, can be detected by scanning the genome for regions showing excess local ancestry from a particular source population relative to the genome average^60^. Applying this approach, we observed two strong deviations in the local ancestry inference with RFMixv2 using European (TSI) as a Middle Eastern proxy, east African (LWK), west African (ESN), and south Asian (STU) populations. One occurs on chromosome 6 at the MHC with 93.5 - 96.5% European ancestry compared to 83% genome-wide (3.2 – 4.5 SD; Supplementary Figure 2). The other, and strongest, deviation in local ancestry is on chromosome 1 in a region encompassing the *ACKR1* gene with 76.5% local east African ancestry compared to 11% genome-wide (24.5 SD; Figure 2A) and corresponds with a drop in European ancestry (23.5%, −19 SD; Figure 2B). This excess African ancestry at *ACKR1* is more extreme than has been reported in other African-admixed populations, both in the AP and elsewhere (Supplementary Table 3).

**Figure 2.**
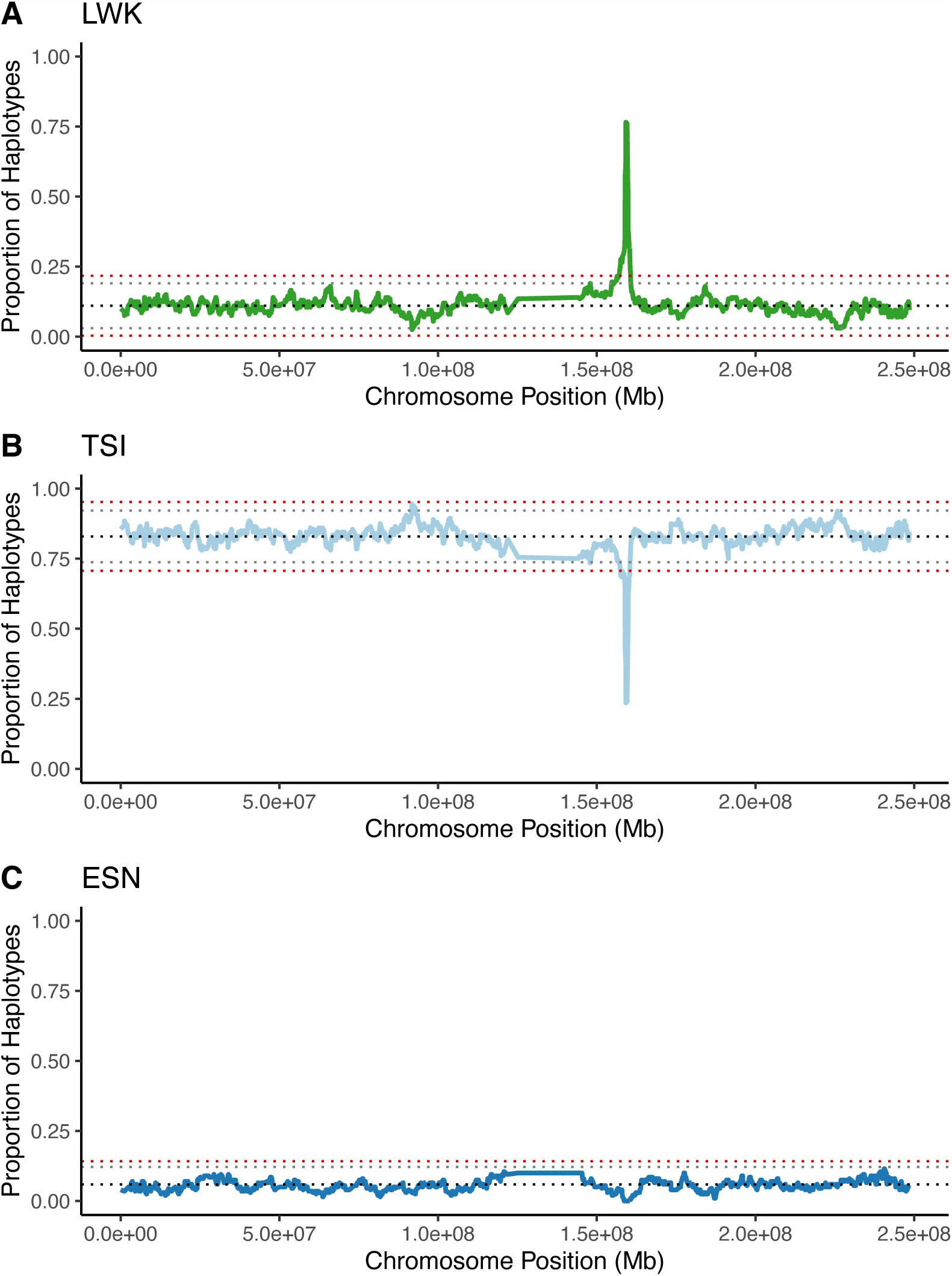
Local ancestry across chromosome 1 in Omanis. In each panel, the genome-wide mean is represented by the dotted black line, three standard deviations (SD) by the dotted gray lines, and 4 SD by the dotted red lines. A) LWK ancestry (East Africa). B) TSI ancestry (Europe). C) ESN ancestry (West Africa).

The Fy^ES^ allele, associated with reduced susceptibility to *P. vivax*, lies within the region of increased east African ancestry (Supplementary Figure 3). We confirmed the frequency of the Fy^ES^ allele in Oman (AF = 0.89) is significantly higher than expected under a neutral model where the expected frequency is a linear combination of the allele frequencies in source populations weighted by their admixture contributions^61^. Lastly, although the Fy^ES^ allele is reported in populations across sub-Saharan Africa, the local ancestry analysis does not infer excess west African ancestry at *ACKR1* (0% ESN ancestry, −2.8 SD; Figure 2C), indicating a likely east African origin for the haplotypes carrying Duffy null alleles in Oman. Together, these suggest strong positive selection for the Fy^ES^ allele post-admixture with east African populations.

In order to quantify the timing and strength of selection for the Fy^ES^ allele, we inferred the selection coefficient using approximate Bayesian computation (ABC)^46^ comparing observed summary statistics to those generated from simulations under a range of parameter values. We used SLiMv4^9,42–45^ to simulate admixture between an east African and ancestral Middle Eastern population. Each simulation sampled from a uniform distribution for initial African ancestry (*m*; range 0.10-0.90) and for the selection coefficient (*s*; range 0.0-0.2). We then used the R package abc^46^ to estimate the selection coefficient based on comparing summary statistics from the simulations to those observed in the Omanis (the proportion of African ancestry at the region encompassing *ACKR1* and variance in African ancestry genome-wide as well as the mean, median and variance in tract length at *ACKR1*). We found that simulations of continuous admixture beginning 10 generations ago (similar to the admixture date inferred by MALDER) was a poor fit for the observed Omani summary statistics, with observed summary statistics falling outside the range of the simulations (Supplementary Figure 4, goodness-of-fit < 0.05), suggesting the Fy^ES^ allele was likely not introduced in a recent pulse of admixture. Models simulating admixture beginning 59 generations ago (matching the older date inferred by MultiWaver) were a much better fit, yielding a selection coefficient estimate of 0.031 (95% CI 0.029-0.034) (Figure 3A), which is slightly lower than has been inferred in other African admixed populations^8,9^. To test whether older admixture, even if smaller in proportion could also be consistent with the observed ancestry at the Duffy locus, we also simulated admixture beginning 75 generations ago, which also fit the observed data, yielding an estimated selection coefficient of 0.036 (95% CI 0.031-0.044) (Figure 3B).

**Figure 3.**
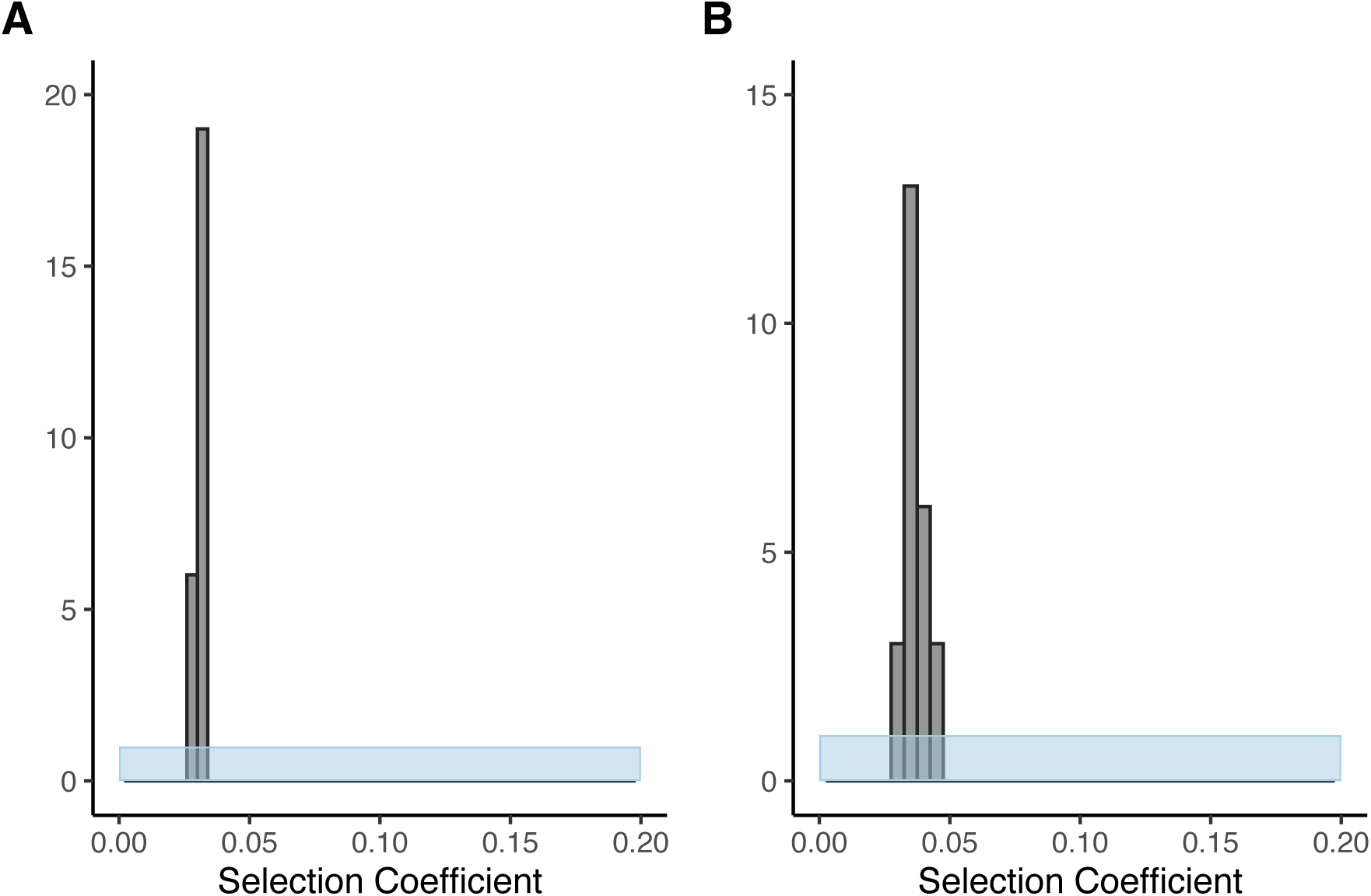
Estimation of selection coefficient with SLiM simulations and approximate Bayesian computation. Each plot shows the prior distribution in blue and the posterior distribution in gray. A) Selection coefficient estimate under a single-pulse model 59 generations ago. B) Selection coefficient estimate under a single-pulse model 75 generations ago.

### Blood group serology confirms high incidence of Duffy negative phenotype and identifies an additional null allele in the Middle East

To further confirm high incidence of the Duffy negative phenotype in Oman, we compared genetic predictions based on the Fy^ES^ SNP (rs2814778; −67 T>C) with Duffy serological phenotypes determined for the 100 Omani donors (collected as part of a panel of blood group phenotypes reported separately, manuscript in prep). Serological absence of both Fy^a^ and Fy^b^ antigens was observed in 87/100 Omani individuals, confirming a high frequency of Duffy negative phenotype. This included all 83 individuals homozygous for the Fy^ES^ allele but also four heterozygous individuals who nonetheless appeared serologically negative for Duffy antigens. Analysis of other variants in *ACKR1* revealed that three of these individuals carried the Duffy X allele (rs34599082; Arg89Cys, known as Fy^X^), associated with reduced Fy^b^ expression^62–64^ that appears below the limit of assay detection.

In the fourth individual, we identified a 2bp frameshift deletion allele (rs773692057; Ser62fs) leading to early protein termination at amino acid 62, thus encoding a loss-of-function allele across all cell types. This allele is observed twice in all of gnomAD^65^ (2/314383 individuals), once in a non-Finnish European individual and once in a Middle Eastern individual. It has also been observed before in the Middle East (frequency of 4/1002 individuals in the Greater Middle East Variome^51^), suggesting it is more common in the Middle East than in other parts of the world. To further assess its relevance in Oman, we considered results from a previous study comparing targeted genotyping to serology in Omani individuals^66^. We identified thirteen individuals similarly predicted by genotype as expressing Fy^b^ antigens but showing a Duffy negative phenotype by serology. We performed Sanger sequencing across the *ACKR1* frameshift position in these additional thirteen samples as well as the frameshift carrier identified here. This validated the presence of the frameshift allele identified in the whole genome sequence data as well as in eight of the additional thirteen individuals, indicating it is indeed more common in Oman.

### Multiple malaria-protective alleles are at higher frequency in Oman compared to other Middle Eastern populations

Given the high frequency of Duffy null alleles in Oman relative to neighboring populations, we aimed to describe patterns of variation across the AP at malaria-associated loci more broadly. To avoid the challenge of comparing variants identified through different analysis pipelines and quality control filters, we obtained raw sequence reads and jointly called variants with Sentieon across the 100 Omanis from this study, 2504 high-coverage individuals from the 1000 Genomes phase 3^67^, 108 Qataris^68^, and 137 individuals from seven Middle Eastern countries (three of which are in the AP)^16^, and here focus on a subset of the genes where variation has been robustly associated with malaria susceptibility: *G6PD*, *HBB*, *GYPA* and *GYPB*.

Glucose-6-phosphate dehydrogenase deficiency (G6PDd), caused by a variety of mutations in the *G6PD* gene, has been associated with protection from both *P. falciparum* and *P. vivax* malarias, though the nature of the association is complex due to its X-linked status and variation in severity of the alleles (class I-IV from most to least severe)^69–71^. We checked the joint call set for any of the missense variants reported in the G6PD mutations database^72^ and identified three in Omanis: A (rs1050829, Asn126Asp, class III, Omani AF=0.04), Mediterranean (rs5030868, Ser188Phe, class II, Omani AF=0.27), and Chatham (rs5030869, Ala335Thr, class II, Omani AF=0.01). These G6PDd alleles are more common in Oman than in the other four AP populations in the joint dataset (Figure 4), with the exception of G6PD Chatham which has a similar frequency in the UAE. We also identified five G6PDd alleles found at low frequencies in Qatari, Yemeni, Emirati, and Saudi populations that are not present in Omanis (Supplementary Table 4).

**Figure 4.**
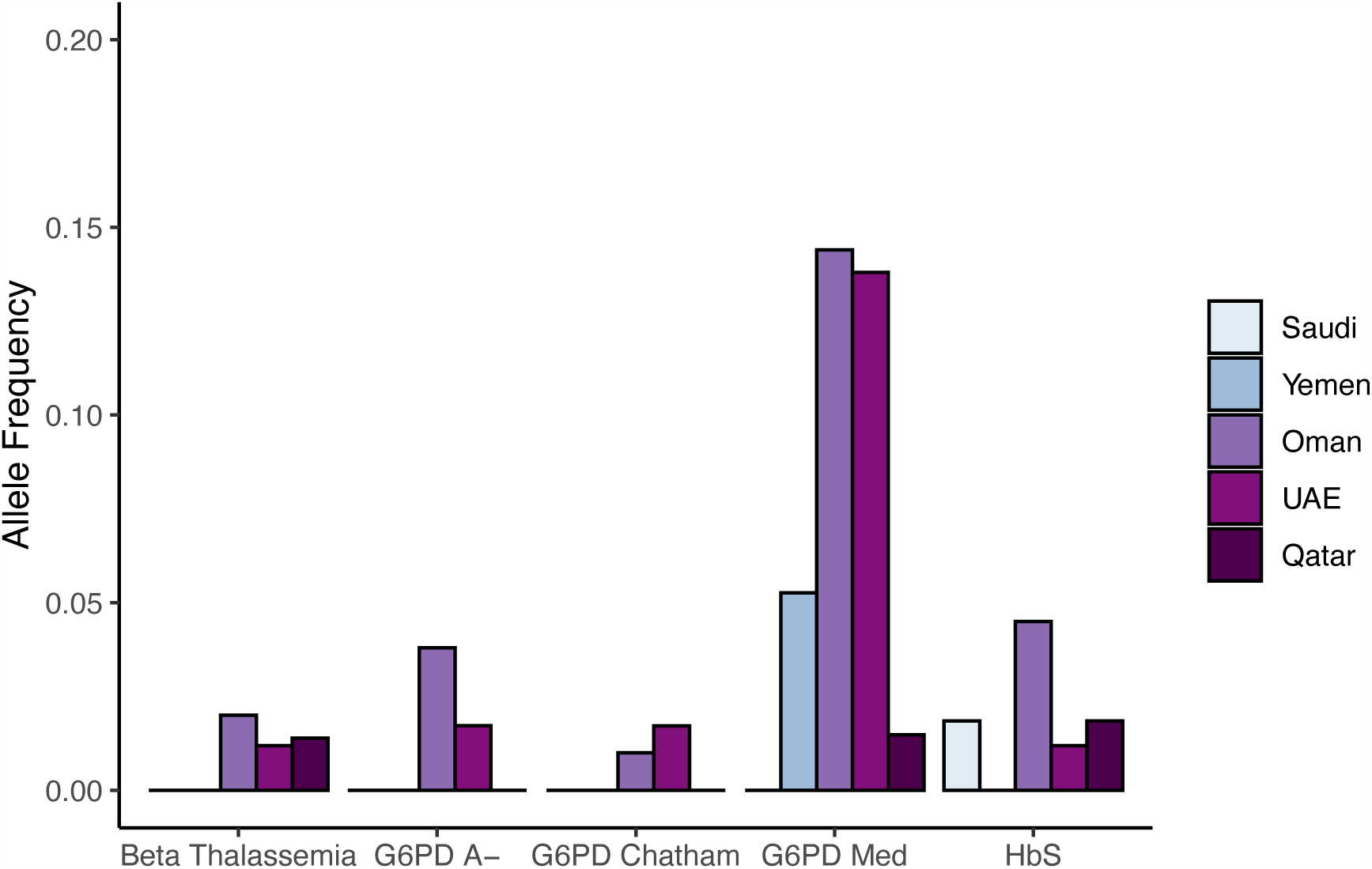
Frequency of select malaria-protective alleles in Omanis and other Arabian Peninsula populations. United Arab Emirates is abbreviated as UAE in the legend.

Multiple alleles affecting the hemoglobin beta gene (*HBB*) have been associated with reduced *P. falciparum* malaria, including both missense changes (such as the sickle cell allele, rs334, Glu6Val) and nonsense alleles (causing *beta*-thalassemias)^73–75^. We found that nine Omani individuals were heterozygous carriers of rs334 (allele frequency (AF) = 0.045), consistent with previously reported frequencies^76,77^. We also identified four Omani individuals heterozygous for distinct *beta*-thalassemia alleles: rs80356820 (Phe46fs), rs33946267 (Glu122Ter), rs35395625 (Pro59fs), and rs33945777 (315+1G>A, splice donor variant). Phe46fs, Glu122Ter, and 315+1G>A have been observed previously in *beta*-thalassemia patients in Oman^78^. Similar to G6PDd alleles, HbS and *beta*-thalassemia alleles are more common in Oman than other AP populations (Figure 4).

Lastly, we investigated copy number variation at the cluster of glycophorin genes that determine the MNS blood group. A complex structural rearrangement affecting both *GYPA* and *GYPB* and encoding the Dantu antigen has been associated with protection from severe *P. falciparum* malaria but is common only in coastal east African populations^47,79,80^. Applying a Hidden Markov Model to infer copy number from coverage data across this locus (including a mappability filter as in Leffler et al. 2017^47^) and looking at tag SNPs, we identified one Omani and one Qatari individual carrying the Dantu structural variant (Supplementary Figure 5; Supplementary Table 5), suggesting it has likely spread to the southern Arabian Peninsula through east African admixture.

Despite higher frequencies than elsewhere in the AP, we did not observe a significant increase in African ancestry at these four malaria-associated loci (average African ancestry = 0.0883, < 1 SD) or a trend towards higher African ancestry at the top 97 loci associated with severe *P. falciparum* malaria identified through GWAS (Supplementary Figure 6).

## Discussion

In this study, we present analyses of whole genome sequence data and blood group serology for 100 healthy Omanis. We find 16% of genetic ancestry in the Omanis is shared with African populations, a majority of which is uniquely shared with the east African LWK population, consistent with previous estimates based on genotyping data^11^. We infer African admixture was introduced into the population via a two-pulse model of admixture with a smaller fraction (0.0725) dating to five generations ago and a larger fraction (0.506) to 59 generations ago. Assuming a generation time of 29 years and a starting date of 1991 (mean birth year of donors), these correspond to pulses around 1846 and 280 CE. The more recent pulse is within the peak period of the Omani empire^81^ and the end of the Indian Ocean slave trade^82^, while the older, larger pulse of admixture is closer to the earliest historical documentation of maritime trade between the regions in the first century AD^56^, predating previous estimates of African admixture into AP populations by 600 years. Previous estimates have ranged from 9-37 generations ago^11,16,53,68,83^, although the older dates in this range were also reported for southern AP populations. Many of the more recent date estimates come from MALDER/ALDER, which we also found to estimate a more recent admixture timing (9 generations), but likely due to the difficulty of LD-based methods to differentiate between multiple pulses from the same source population^40,84^. The only populations in which a two-pulse model of African admixture has been inferred are the Emiratis, Saudis, and Omanis, and only when using a tract length-based approach^11,83^. The older admixture date overlaps with some estimates for the Bantu Expansion into East Africa^85,86^, which could indicate that the present-day Luhya are not representative of the east African population at the time of the inferred pulse 1,800 years ago. However, the local increase and higher global similarity to the LWK population rather than other Bantu groups from central and west Africa suggest it is largely not the case.

Using both sequencing and serological data, we also investigated the frequency and admixture history at the *ACKR1* gene, which has been associated with *P. vivax* susceptibility. We report a high frequency of both the Fy^ES^ allele (0.89) and Duffy negative serological phenotype (87/100). Most Duffy negative individuals are homozygous for the Fy^ES^ allele (96%), where we find slightly but significantly more homozygous genotypes than expected under Hardy Weinberg Equilibrium (0.83 genotype frequency vs. 0.79 expected, *p*-value = 0.0016), possibly reflecting sub-structure with respect to ancestry within the population, as has been suggested in African Americans^87^. The remainder of Duffy negative individuals were heterozygous for the Fy^ES^ allele and either the Fy^X^ allele or a frameshift allele, both reported in Oman here for the first time. Fy^X^ is characterized by low Fy^B^ expression^63,64^, apparently below the limit of detection by our assay when phenotyping compound heterozygotes for the Fy^ES^ allele. Reduction in expression of Fy antigens may provide some protection from *P. vivax* infection^88^. Compound heterozygosity for Fy^ES^ and the true null frameshift allele would result in a lack of DARC on red blood cells, while retaining some expression on endothelial cells. This frameshift allele is extremely rare worldwide but has been reported in several Middle Eastern populations and is documented in the ISBT database of blood group alleles^89^. Here, we found that it explained 60% of individuals with a Duffy blood group mismatch from a recent study comparing serology and targeted genotyping in Oman^66^, indicating it should be incorporated in future targeted genotyping approaches for blood group inference in Oman and other AP populations. Whether it is associated with any disease phenotype in homozygotes is unknown.

The high frequency of the Fy^ES^ allele in Oman is accompanied by a striking signal of increased east African ancestry. In the region encompassing the *ACKR1* gene, 76% of haplotypes are inferred to have local east African ancestry compared with the genome-wide mean of 11%, suggesting positive selection following introduction of the Fy^ES^ allele into the Omani population. This is slightly more than has previously been inferred based on genotype data in Oman (increase from 11% to 49%)^11^ but similar to what has been reported based on sequencing data in Yemen (9% to 74%)^16^, possibly reflecting higher sensitivity of ancestry inference based on sequencing data, and a similar admixture and selection history in Oman and Yemen.

Although the larger increase in African ancestry at *ACKR1* in Yemen and Oman relative to other admixed populations could be due to stronger selection, we find rather that it is consistent with a slightly lower selection coefficient but an older introduction. Simulations of more recent introductions do not match observed summary statistics for tract length distribution, whereas simulations under models from 50-75 generations match well. While little documentation exists regarding the presence of *P. vivax* in Oman beyond the early 1900s, there is evidence of agriculture in some regions of Oman dating back to the first millennium AD^90^. Agriculture has been linked to malaria transmission in several countries^91–93^, and *P. vivax* may have been present in Oman by the time Fy^ES^ was introduced. Whether this could have driven strong selection, however, is unclear. Similar selection at *ACKR1* has now been observed convergently across multiple populations in the Middle East (Supplementary Table 3), which overall has been arid for the last 20,000 years^94^ and much of the peninsula does not currently support malaria transmission, indicating only partial malaria endemicity at most^25^. The accumulating evidence for repeated, convergent selection with a similar selection coefficient across populations with very different ecological settings also raises the possibility that another selective pressure could be involved. Moreover, a signal of significant reduction in European ancestry at *ACKR1* was observed in the Fulani population in west Africa after recent European admixture, suggesting ongoing selection for the Fy^ES^ allele despite very low prevalence of *P. vivax*^95^.

The Fy^ES^ allele has been associated with diseases and pathogens aside from malaria, including *Staphylococcus aureus*, a bacterium which causes several illnesses including sepsis, pneumonia, osteomyelitis, skin infections, and endocarditis^96^. DARC also serves as a receptor for hemolytic leukocidins, toxins produced by *S. aureus* facilitating immune evasion, proliferation, and virulence^96^. DARC expression has also been associated with lung^97^ and breast cancer metastasis and disease progression^98–100^, likely due to the role of DARC in immune response and chemokine regulation. Lastly, the Fy^ES^ allele is associated with low neutrophil counts, but no health risks or benefits have been identified^87^. It is therefore possible that an alternative, or additional, selective pressure may be contributing to the adaptive admixture signals at the Fy^ES^ allele. In any case, the high frequency of Fy^ES^ and presence of other alleles with low or no Duffy expression have likely led to lower transmission of *P. vivax* in the southern AP, much as it is thought to have excluded *P. vivax* to very low levels in sub-Saharan Africa.

In contrast, we do not find signals of higher local African ancestry at *P. falciparum*-associated loci. This is perhaps not surprising given the more polygenic architecture, fitness costs limiting frequency of some alleles, and allelic heterogeneity documented across populations including outside Africa. For example, the HbS allele is found on at least five different haplotype backgrounds, one of which is more common in the Middle East and India (Arab-Indian haplotype)^101^. In the Omanis sequenced here, the HbS allele is found on the Arab-Indian haplotype in three of the nine carriers, none of which were inferred to have local African ancestry at the locus. There are numerous different SNVs that cause G6PD deficiency found in populations throughout the world^72^ as well as beta-thalassemias, which are also at relatively low frequency potentially due to balancing selection. We identify the *P. falciparum*-associated Dantu structural variant (SV), which has only been reported at appreciable frequency in coastal East Africa^47,79,80^. However, it was carried by a single Omani individual who had a high proportion of African ancestry genome-wide, suggesting it may result from more recent admixture.

Our findings support the repeated convergent selection of Fy^ES^ in diverse African admixed populations with remarkably similar strength and pattern of selection, despite different timing and geographical and ecological milieus. The signature of adaptive admixture is particularly striking in Oman, which we find to be due to an earlier introduction with lower overall African ancestry than other populations in which this signal has been reported. Our identification of two additional alleles that contributed to the Duffy negative phenotype in Oman alludes to the possibility that alleles could be present in other populations where the Duffy negative phenotype has been under selection. The resulting high frequency of Duffy negative phenotypes in Oman has likely maintained a low frequency of *P. vivax* in the last century, leading to a higher proportion of malaria cases due to *P. falciparum*. However, it remains unknown if *P. vivax* could have sustained a similar enough selective pressure across multiple human populations and geographic locations to observe this unique signature of adaptive admixture which has occurred at various time points in the last 2,000 years.

## Declaration of Interests

G.B.J.B. is an employee of Allelica, Inc.

## Acknowledgments

We thank all blood donors for participation in this study. We would like to thank David Bresnahan, PhD from the University of Utah Department of History for information about key historical events along the Swahili coast and across the Indian Ocean, Alan Rogers, PhD from the Department of Anthropology for feedback on the manuscript, and the Leffler lab for discussion. This work was supported by the Omani Ministry of Higher Education, Innovation and Research (formally known as the Research Council, grant number RC/RG-MED/HAEM/18/02) and by the United States National Institutes of Health (grant number R35 GM147709 to E.M.L.). The support and resources from the Center for High Performance Computing at the University of Utah are gratefully acknowledged. The computational resources used were partially funded by the NIH Shared Instrumentation Grant 1S10OD021644-01A1. P.E.H. was supported by T32 GM007464 and T32 GM141848.

## Author Contributions

Conceptualization – A.Z.A., G.B.J.B., and E.M.L.; Methodology – P.E.H., S.A., A.A.S., and E.M.L.; Validation – E.M.L.; Formal Analysis – P.E.H.; Investigation – P.E.H., M.A., and S.A.H.; Resources – A.Z.A., S.A. and S.A.M.; Writing – Original Draft, P.E.H., A.Z.A., and E.M.L.; Writing – Review & Editing, A.Z.A. and G.B.J.B; Visualization – P.E.H.; Supervision – A.Z.A., S.A., A.A.M., and E.M.L.; Project Administration – A.Z.A.; Funding Acquisition – A.Z.A. and E.M.L. All authors have read and approved this manuscript.

## Data and code availability

Short read sequences and variant calls generated in this study have been deposited in dbGaP (Accession Number XXX) with controlled access following the terms of consent provided to study participants.

## Notes

### Competing Interest Statement

G.B.J.B. is an employee of Allelica, Inc. All other authors declare no competing interests.

