## Supplemental Figures and Tables for "Adaptive admixture at *ACKR1* (the Duffy locus) may have shaped *Plasmodium vivax* prevalence in Oman"

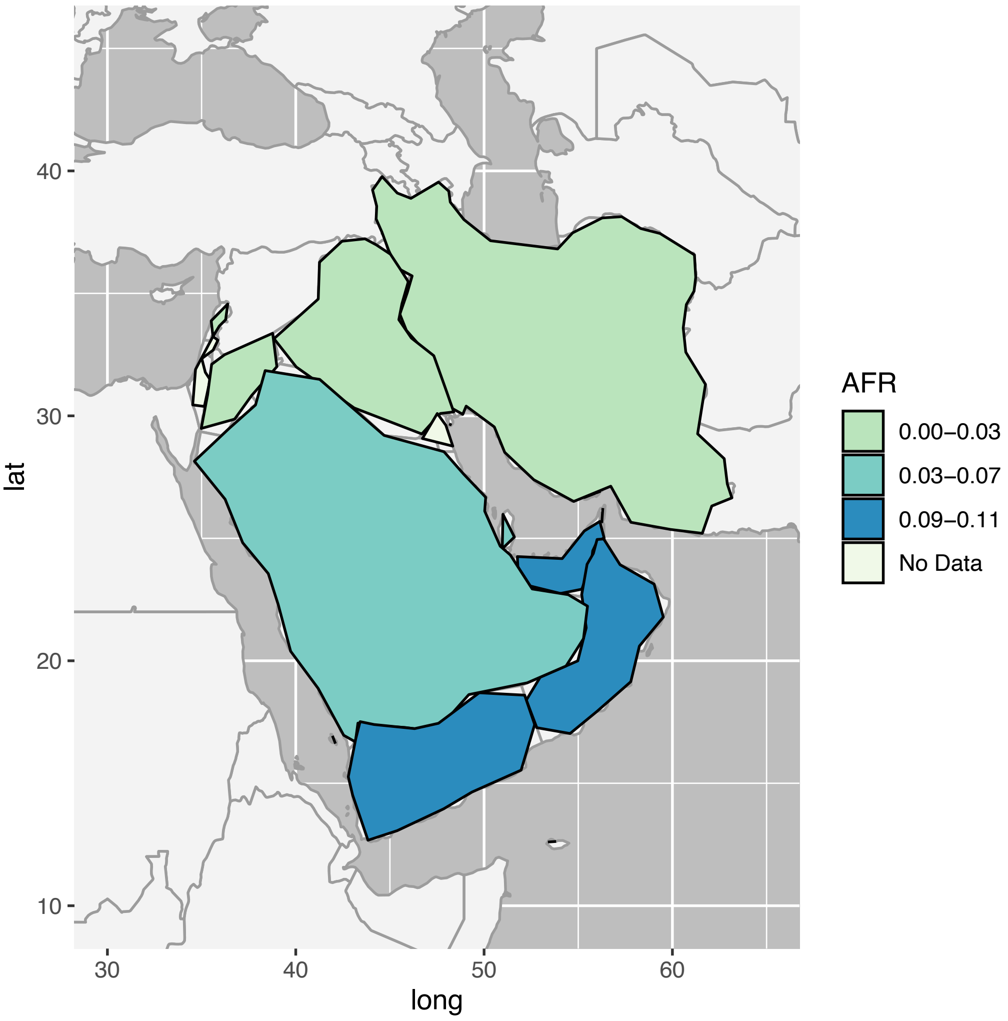


**Figure S1.** Distribution of inferred genome-wide African ancestry in Arabian Peninsula populations from Fernandes et al. 2019[1] and Almarri et al. 2021[2]**.** The plot was produced with the `maps` R package.

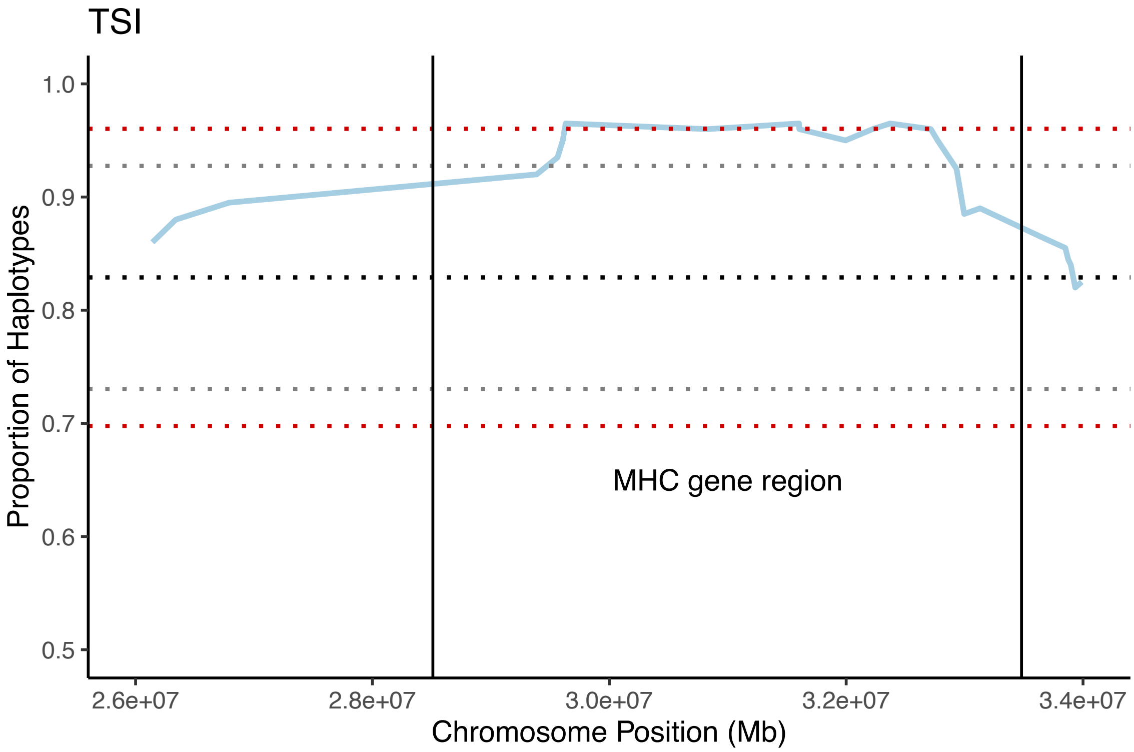


**Figure S2.** TSI (European) local ancestry inference along chromosome 6 across the MHC region. The dotted black line represents the genome-wide mean (0.829) and the dotted gray and red lines represent three and four standard deviations (SD) from the mean, respectively (1 SD = 0.0328).

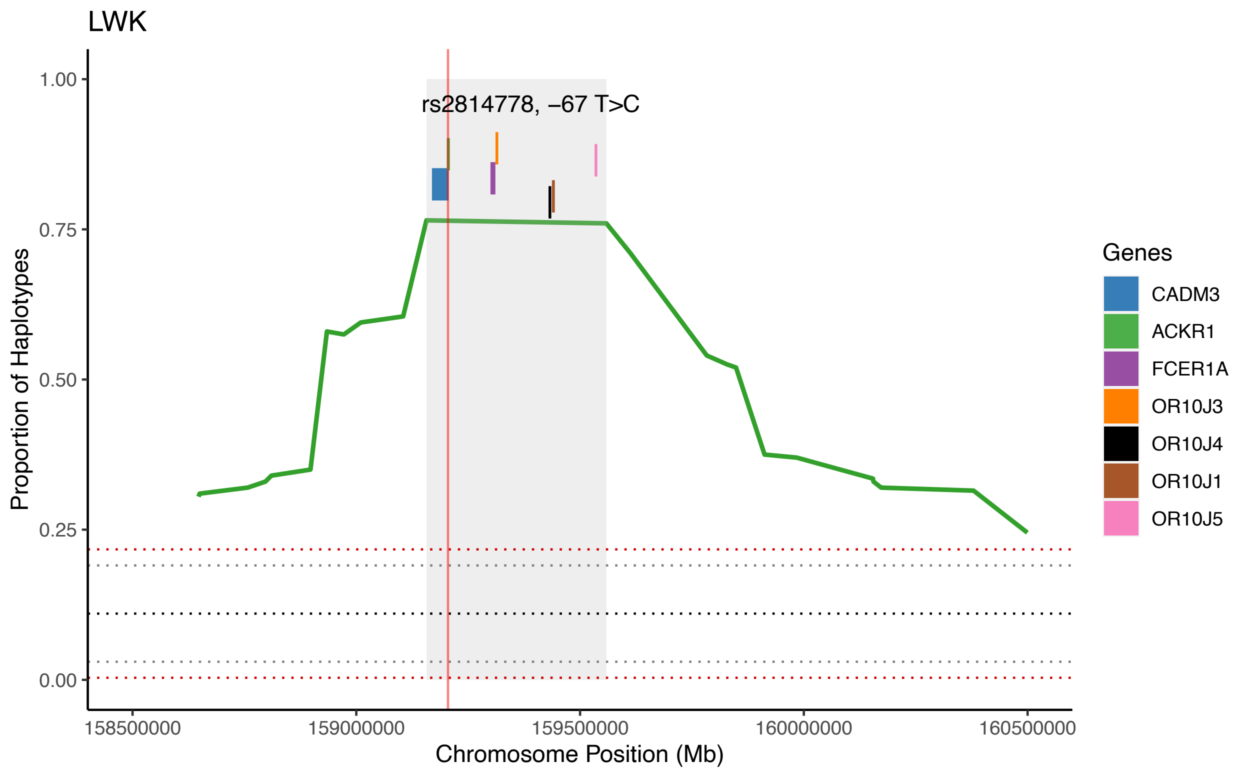


**Figure S3.** LWK (east African) local ancestry inference along chromosome 1 across the *ACKR1* region. The dotted black line represents the genome-wide mean (0.11) and the dotted gray and red lines represent three and four standard deviations (SD) from the mean, respectively (1 SD = 0.0267). The shaded gray region covers the locus with the highest proportion of LWK ancestry, and the genes in the region are indicated by colored rectangles. The $Fy^{ES}$ SNP is marked by the vertical red line.

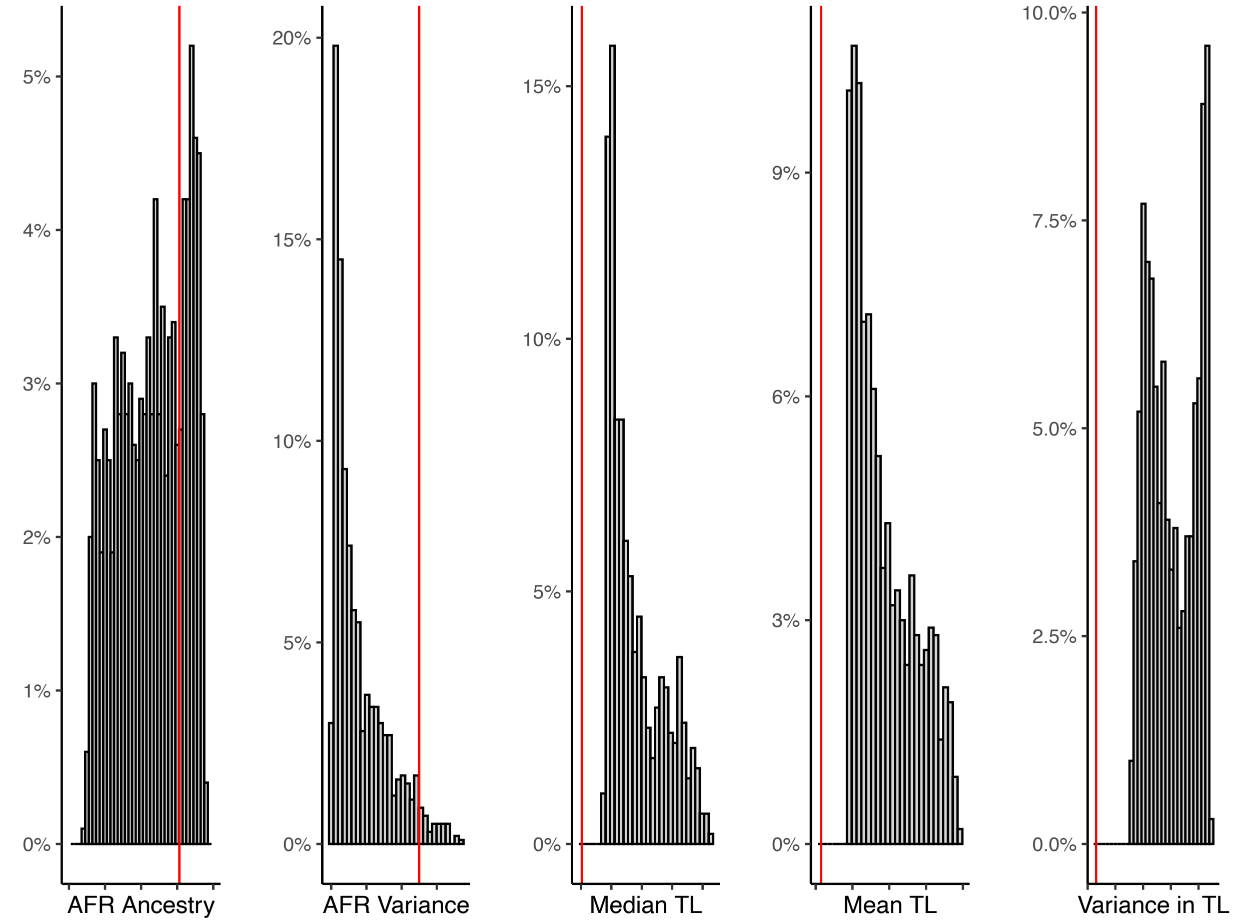


**Figure S4.** Comparison of summary statistics for simulations at 10 generations and observed statistics in the Omanis. In each plot, the vertical red line represents the value observed in the Omanis and the histogram is the distribution of expected values. From left to right, the plots are the proportion of African (AFR) ancestry at *ACKR1*, variance in global AFR ancestry, median tract length (TL) at *ACKR1*, mean TL at *ACKR1*, and the variance in TLs at *ACKR1*.

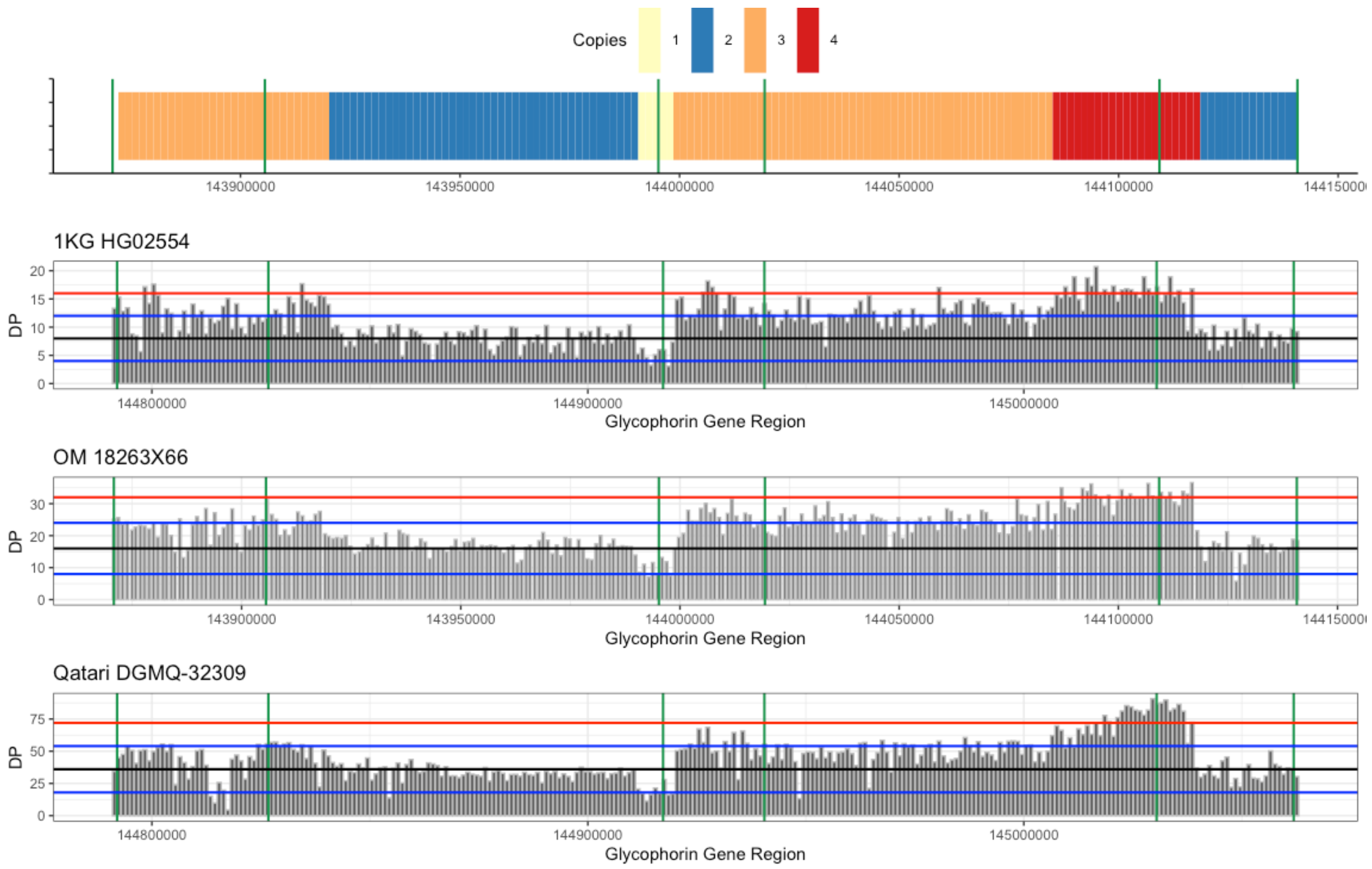


**Figure S5.** Read depth (DP) across the glycophorin gene region in 1kb bins in the individual carrying the Dantu structural variant from the 1000 Genomes dataset (HG02554) and the two carriers inferred here from Oman and Qatar. The top plot represents the copy number inference for HG02554. HG02554 and Qatari DGMQ-32309 are aligned to hg19 whereas the OM 18263X66 sample is aligned to GRCh38. In the bottom three plots, the black solid line is the genome-wide coverage, blue lines are 1.5 or 0.5 times the mean coverage, and the red line is 2 times the genome-wide coverage. The vertical green bars from left to right delineate the GYPE, GYPB, and GYPA gene boundaries in all plots.

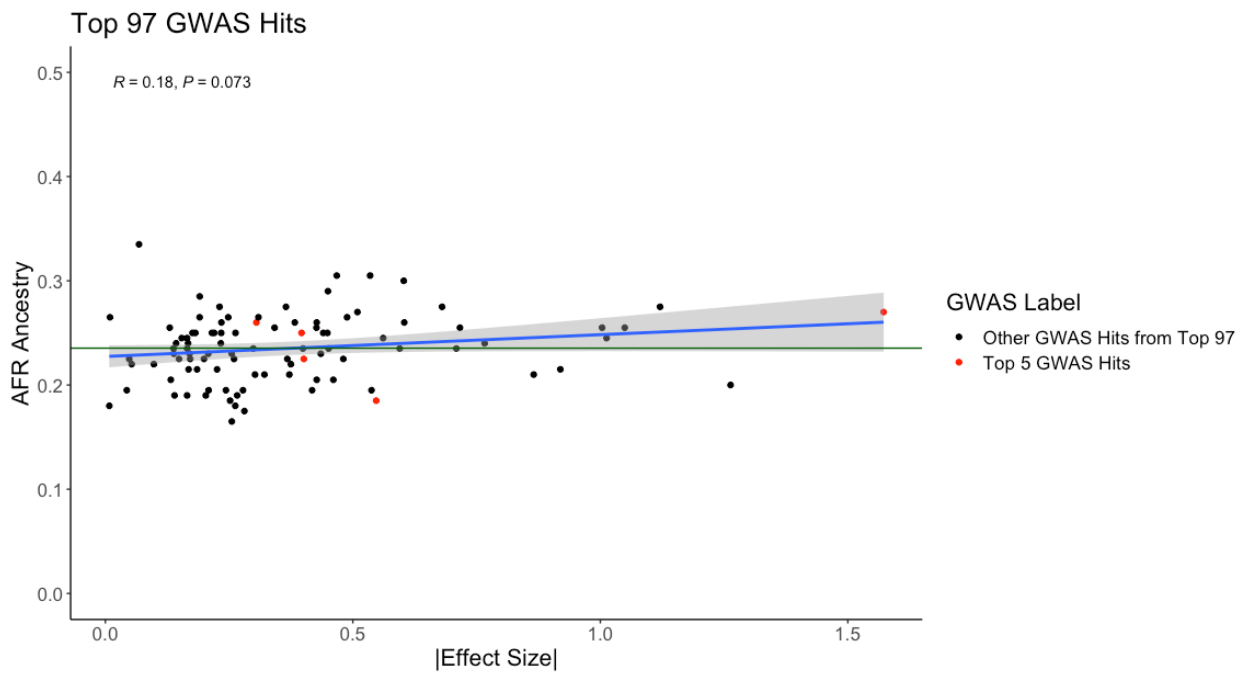


**Figure S6.** Absolute value of the effect size of the 97 most associated variants from GWAS of severe *P. falciparum* malaria[3] compared to the proportion of east African ancestry inferred at each variant. The horizontal line represents the mean genome-wide African ancestry in Oman. The blue line represents the correlation between African ancestry and effect size, with a 95% confidence interval represented by the shaded gray region.

**Supplemental Tables**

**Table S1.** Timeline of key events between Oman and East Africa in documented history. Information is compiled from: Casson 1989[4], Al-Mahdhouri 2014[5], Al-Maamiry 1988[6], Al-Maamari 1992[7], and Al-Riyami 2016[8].

| Time Frame | Event |
|---|---|
| 0 – 100 CE | First documented evidence of trade networks between East African and Omanis as well as other networks across the Indian Ocean |
| 600 – 700 CE | Omanis migrated to the Lamu Archipelago off the coast of Kenya |
| 1200 – 1300 CE | Omanis migrated to Bathi in southwest Kenya |
| 1600 – 1700 CE | Omanis migrated to Mombasa on the coast of Kenya |
| 1696 – 1698 CE | Siege of Mombasa and beginning of the Omani Empire |
| 1696 – 1856 CE | Reign of the Omani Empire including parts of Kenya, Tanzania, Southern Iran, and Pakistan |
| 1832 – 1856 CE | Zanzibar is the capital of Oman |
| 1872 CE | Treaty ending Indian Ocean Slave Trade in Oman |
| 1964 CE | Zanzibar revolution leading to the reverse migration of Omanis from Africa to Oman |

**Table S2**. The number of SNPs, INDELs, and total variants called in 100 Omani samples. The second column represents the intersection with 1000 genomes, merged by position and allele.

| | Count | Merged Count |
|---|---|---|
| **SNPs** | 22,039,817 | 17,275,800 |
| **INDELs** | 3,933,262 | 1,337,279 |
| **Total** | 25,778,021 | 18,601,784 |

**Table S3.** Local African ancestry genome-wide and at the Duffy locus in published studies of African-admixed populations. If available, the date of African admixture in generations and the selection coefficient at the Duffy locus are included.

| **Admixed Population** | **Reference** | **Sample Size** | **Data Type** | **Number of SNPs** | **Average African Ancestry (%)** | **African Ancestry at *ACKR1* (%)** | **Date of Admixture (gen)** | **Selection coefficient** |
|---|---|---|---|---|---|---|---|---|
| Malagasy | Pierron et al. 2018. Nat Comm. | 700 | SNP Array | 1.9e+06 | 62% | 91% | 27 | >0.2 |
| Santiago | Hamid et al. 2021. eLife. | 172 | SNP Array | 8.8e+05 | 73% | 83% | 20 | 0.08 |
| Makranis | Laso-Jadart et al. 2017. AJHG. | 24 | SNP Array | 1.3e+06 | 22% | 44% | 12 | NA |
| Saudi Arabians | Fernandes et al. 2019. Mol Biol Evol. | 100 | SNP Array | 1.7e+05 | 4-7% | 28% | 2 and 31 | NA |
| Yemenis | Fernandes et al. 2019. Mol Biol Evol. | 100 | SNP Array | 1.7e+05 | 11% | 61% | 27 | NA |
| Omanis | Fernandes et al. 2019. Mol Biol Evol. | 100 | SNP Array | 1.7e+05 | 11% | 49% | 5 and 34 | NA |
| Emiratis | Fernandes et al. 2019. Mol Biol Evol. | 120 | SNP Array | 1.7e+05 | 11% | 35% | 5 and 22 | NA |
| Iranians | Fernandes et al. 2019. Mol Biol Evol. | 80 | SNP Array | 1.7e+05 | 0% | 16% | NA | NA |

**Table S4**. Glucose-6-phosphate dehydrogenase deficiency (G6PDd) alleles from the G6PD mutation database[12] found in Arabian Peninsula populations, but not in Omanis.

| rsID | Amino Acid Change | Populations (# Indv.) | Mutation Name | Class |
|---|---|---|---|---|
| rs76645461 | Ile48Thr | Qatar (1), Saudi Arabia (1) | Aures | III |
| rs1050828 | Val68Met | Qatar (2), United Arab Emirates (2) | Asahi | III |
| rs370918918 | Met159Ile | Qatar (1) | Gond | NR |
| rs5030872 | Asp181Val | Saudi (1) | Malaga | III |
| rs72554664 | Arg463His | Yemen (1) | Kaiping | II |

**Table S5.** Genotypes of marker SNPs in highest LD with the Dantu structural variant in the HG02554 sample known to carry Dantu as well as the Omani and Qatari samples inferred to carry Dantu by read coverage. Data is from the joint calling dataset including the high coverage 1000 genomes, 108 Qatari samples, and 137 Middle Eastern samples.

| | AC | AN | HG02554 | OM 18263X66 | Q DGMG-32309 |
|---|---|---|---|---|---|
| 4:143653351 | 3 | 5698 | 0/1 | 0/1 | 0/1 |
| 4:143638460 | 3 | 5698 | 0/1 | 0/1 | 0/1 |
| 4:143639385 | 3 | 5698 | 0/1 | 0/1 | 0/1 |
| 4:143664382 | 3 | 5698 | 0/1 | 0/1 | 0/1 |
| 4:143554460 | 3 | 5698 | 0/1 | 0/1 | 0/1 |
